# Episodic-Like Memory in a Simulation of Cuttlefish Behavior *

**DOI:** 10.1101/2025.09.03.674043

**Authors:** Sriskandha Kandimalla, Qian Y. Wong, Kary Zheng, Jeffrey L. Krichmar

**Affiliations:** Department of Cognitive Sciences, University of California, Irvine, Irvine, CA 92697-5100; Department of Psychology, University of California, Irvine, Irvine, CA 92697-7085; Department of Cognitive Sciences, Department of Computer Science, University of California, Irvine, Irvine, CA 92697-5100

## Abstract

Episodic memory involves remembering the what, when, and where components of an event. It has been observed in humans, other vertebrates, and cephalopods. In clever behavioral experiments, cuttlefish have been shown to have episodic-like memory, where they demonstrate the ability to remember when and where a preferred food source will appear. The present work replicates this behavior with a parsimonious model of episodic memory. To further test this model and explore episodic-like memory, we introduce a predator-prey scenario in which the agent must remember what creatures (e.g. predator, desirable prey, or less desirable prey) appear at a given time and region of the model environment. This simulates similar situations that cephalopods face in the wild. They will typically hide when predators are in the area, and hunt for prey when available. When the memory model is queried for an action (e.g., hunt or hide), the cuttlefish agent hunts for preferred food, like shrimp, when available, and hides at other times when a predator appears. When the memory model is queried for a place, the cuttlefish agent acts opportunistically, seeking less-preferred food (e.g., crabs) if it is located farther from a predator. These differences show how behavior can be altered depending on how memory is accessed. Querying the model over time might mimic mental time travel, a hallmark of episodic memory. Although developed with cephalopods in mind, the model shares similarities with the hippocampal indexing theory and captures aspects of vertebrate episodic memory. This suggests that the underlying mechanisms supporting episodic-like behavior in the present model may not be unique to cephalopods.

## 1 Introduction

Episodic memory involves the recollection of a personal experience. Recalling these experiences requires mental time travel to reflect on the past [38]. Endel Tulving, a pioneer in episodic memory research, suggests that this is a uniquely human capability [38]. Because we don’t speak the same language as non-human animals, it is difficult to ask these organisms to recall a memory and place it in time. Such recall would require a notion of self-awareness and conscious recollection. However, other animals have demonstrated episodic-like memory behaviorally by recalling the “what”, “when”, and “where” of past events [8]. Because directly querying declarative memory in non-human animals is challenging, whether they possess *episodic memory* remains an open question. Hence, the term episodic-like memory is used in much of the animal literature to describe the recollection of events.

It has been suggested that episodic memory requires a hippocampus [39]. The discovery of rodent hippocampal place cells in the early 1970s demonstrated that there was a neural correlate of “where” in the rodent brain [25]. Later studies showed recall of place cell sequences during sleep and quiet waking [27, 42]. Furthermore, place cells show both retrospective coding of past locations and prospective coding of future locations [11]. This suggests something like a “when” neural correlate in the hippocampus.

Episodic-like memory has been demonstrated in non-mammalian species that contain a hippocampus. For example, it has been shown that scrub jays remember when food items are stored by allowing them to recover preferred perishable food and non-perishable food [7]. Scrub jays searched preferentially for fresh food if not much time passed. But if enough time had passed that the preferred food had decayed, they searched for the non-perishable food. These behavioral experiments demonstrated that corvids could recall the “what” (food type), “when” (short vs. long delay), and “where” (cache location) of their memory. Although birds have a hippocampus, the avian brain lacks a cortex and the highly processed multimodal hippocampal inputs that are observed in the mammalian brain [8].

Cuttlefish, invertebrates in the same class of cephalopods as octopus, have shown episodic-like memory in experiments similar to those carried out with scrub jays [17, 31]. In the first phase of these experiments, cuttlefish were trained to remember the locations of a preferred food (shrimp) and a non-preferred food (crabs). In the second phase of the experiment, cuttlefish learned that the preferred food was only available after a long delay, and were able to hold off feeding on the non-preferred food until the preferred food was presented. Like the scrub jays, cuttlefish recalled that shrimp (what), were available after a delay (when), at a specific location (where).

Interestingly, these invertebrates do not have a hippocampus, rather learning is thought to take place in their vertical lobe [14, 24]. The anatomy of the vertical lobe has similarities to the hippocampus [33]. There are strong fan-in signals from visual and chemotactile regions, and fan-out signals to motor areas. The neurons in this brain area show signs of long-term plasticity [32]. Such architecture may support episodic-like memory, as suggested by both experimental and theoretical work.

The hippocampal indexing theory, proposed by [35, 36], supports such an architecture. The idea is that the mammalian hippocampus does not contain the memory itself, rather it has pointers to the cortex to form and retrieve memories. The multimodal information from the cortex is converted to an index that activates a set of neurons in the hippocampus. If a new memory is experienced, a new hippocampal index code is generated and the connections from the activated hippocampal neurons back to the cortical columns associated with the memory are strengthened. If a memory is to be recalled, a set of cortical columns are input to the hippocampus and the memory is read out. Furthermore, it has been shown that hippocampal neurons in humans encode conjunctions of a memory [19]. Similarly, the vertical lobe could take multiple inputs encoding “what”, “when”, and “where” to encode the appropriate memory or action. In computer science, this is like a content-addressable memory or a database that can be queried [15, 21].

We constructed a model loosely based on the hippocampal indexing theory to demonstrate episode-like memory. The memory model is a three-dimensional matrix, indexed by what, when, and where tuples, which can also be queried along individual dimensions. This memory architecture is sufficient to replicate cuttlefish episodic-like memory experiments in simulation. To further challenge the model, we created a predator-prey scenario in which the simulated cuttlefish agent had to remember what item (i.e., predator, crab, shrimp) would appear when, and where could it find prey and avoid predators. Depending on how the memory was queried shaped the agent’s behavior. Whereas asking what object can be found at the current time led to delaying hunting, asking where the agent should go at this time led to opportunistic hunting of a safer, non-preferred food. Multiple queries of the memory can result in mental-like time travel. These results show how a simple architecture might support episodic-like behavior. We will discuss how this could be expanded in future iterations.

## 2 Materials and Methods

Two episodic memory scenarios were developed. The first modeled experiments that demonstrated episodic-like memory in cuttlefish [17]. The second involved predator and prey, and was based on cuttlefish behavior in the wild. Like [28], the agent had to distinguish between predators and prey, and then take appropriate action. The present scenario requires memory for when and where these events occur. The source code for both these simulations is written in Python and publicly available at: https://github.com/jkrichma/EpisodicLikeMemoryModel.git

### 2.1 Episodic Memory Model

At the core of both scenarios is a three-dimensional memory model that is indexed by “what”, “when”, or “where”, as well as combinations of these indices (Figure 1). The structure holds the expected values of these indices. For example, a query of “what[shrimp]” and “when[hour 3]” would return the expected values of shrimp at the 3^rd^ hour over all locations in the environment (see Equation 1).

**Figure 1.**
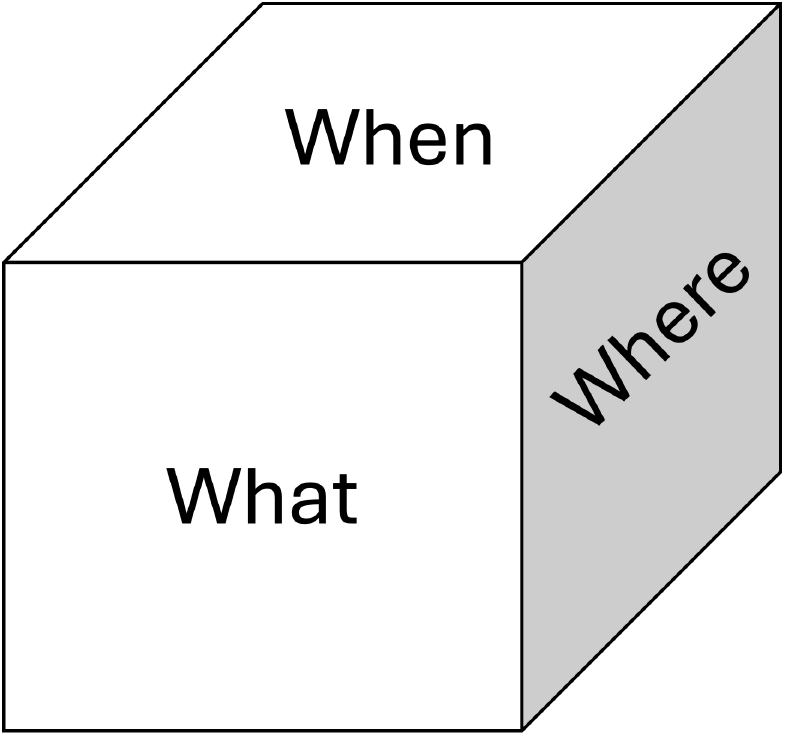
Episodic memory model. Three-dimensional structure that can be indexed in one or more dimensions. For example, a query of “What” and “When” returns the expected values of an item at a given time over all possible locations “Where”.

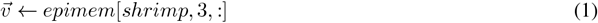

The result from this query is the vector 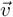 that contains the expected value of shrimp at all the “where” locations in memory. This vector can then be used by a reinforcement learning algorithm to select the appropriate action (e.g., hunting or roaming), with an exact action set depending on the scenario described below.

We use the common *delta rule* to learn associations of what, when, and where. More sophisticated learning rules could be applied but the delta rule will suffice for these simulations to demonstrate episodic-like memory. The learning rule is given by Equation 2.

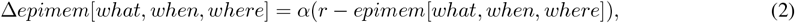

where *α* is the learning rate and *r* is the reward or penalty incurred.

For example, if a shrimp was found and eaten at the 3^rd^ hour at location 42 and received a reward of 4 points with learning rate of 0.10, then Equation 3 would be:

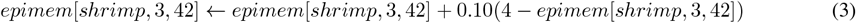

Once a value vector [ineq is acquired for a given action, we subject it to the Softmax function. In the case of equation 1, the vector would be a list of expected values at each location and the action *act* given by Equation 4 would be go to a specific location.

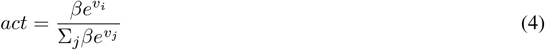

Combinations of memory queries can be used to obtain expected values and agent actions. For example, one could query “what” and “when” for each object (e.g., crab, shrimp, predator), then subject these values to equation 4, which may choose *hunt shrimp* as the best action at this time. Then an additional query of “shrimp” and “current time” would give the expected location of shrimp at the present time.

### 2.2 Simulation Environments

We describe two different scenario environments to test the episodic memory model. In both cases, we used a grid world containing a cuttlefish agent, shrimp, crabs, predators. Predators were assumed to detect the cuttlefish at a greater distance, while the cuttlefish’s vision allowed it to only perceive nearby objects.

#### 2.2.1 Episodic Like Memory Scenario

In [17], episodic memory in cuttlefish was investigated in a two phase experiment. In the first phase, the cuttlefish learned where crab and shrimp were located. In the second phase, the cuttlefish learned that shrimp, which is a preferred food, was only available after a 3 hour delay. To simulate this experiment, an environment was created with two objects, a crab and a shrimp, a 3 hour duration, and an 8×8 grid world environment. This created an episodic memory that was initialized to:

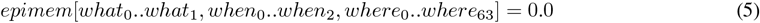

As in [17], both the crab and shrimp are available during all 3 hours in the first phase of the experiment. If the agent reaches one of these food items, the appropriate reward (see Table 1) is applied to the memory as given by Equation 2. In the second phase, the crab is always available, but the shrimp location is only rewarded after 3 hours.

**Table 1.** Episodic Like Memory Scenario Parameters.

|  |  |
| --- | --- |
| $\alpha$ | 0.10 |
| $\beta$ | 1.0 |
| Cuttlefish Initial Position (x,y) | (0, 4) |
| Cuttlefish Vision | < 2 |
| Crab Reward | 1.0 |
| Crab Position (x,y) | (7, 0) |
| Shrimp Reward | 4.0 |
| Shrimp Position (x,y) | (7, 7) |
| Roam Value | 0.5 |

The simulation ran for 100 days, where each day lasted 3 hours, and an hour lasted 100 simulation time steps. The three actions were: *hunt crab, hunt shrimp*, and *roam*. The value for *roam* was set to 0.5, the other action values were learned by the model (see Equation 2). At the start of every hour: 1) Values were acquired with query for each object (crab, shrimp) at the current time (see Equation 1). The maximum value across all locations was chosen. 2) An action was chosen with the Softmax function (see Equation 4). The average value or a sum of values could be used, as will be seen in the other scenario. 3) If the action is to *hunt*, an additional query then retrieves the location where the chosen object has its maximum value. It is assumed that the cuttlefish agent knows how to reach this location. 4) If the action is to *roam*, then the cuttlefish agent moves randomly a step in one of 8 cardinal directions. 5) For all 3 actions, if a shrimp or crab is available at this time, and the cuttlefish agent is less than 2 steps from the object, the value is updated by Equation 2. The parameters for this scenario are given in Table 1.

#### 2.2.2 Predator-Prey Scenario

To further test our episodic memory model, we created a predator-prey scenario in which the cuttlefish agent learned when and where shrimp and crab were likely to be located, but also needed to learn when and where a predator could be located. To simulate this experiment, an environment was created with three objects, a crab, a shrimp, and a predator, a 6 hour duration, and a 12×12 grid (see Figure 3). This created an episodic memory that was initialized to:

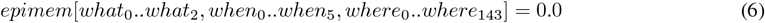

The simulation ran 200 days, where each day lasted 6 hours, and an hour lasted 100 simulation time steps. The four actions were: *hunt crab, hunt shrimp, hide*, and *roam*. The value for *roam* was set to 0.5, the other action values were learned by the model (see Equation 2). At the start of each hour, objects were placed in the appropriate regions of the environment (see Figure 3). If the action was *hunt*, the cuttlefish agent headed to the center of region where the prey was most likely to be found. Once in the region, the agent randomly roamed the region until a prey was found, whereby it receives the appropriate reward, or the hour had passed. If the action was *hide*, the agent simulated camouflaging. That is, the cuttlefish was invisible to the predator. The predator randomly roamed the environment until it saw the cuttlefish was within a distance of 4. In that case, the predator would head directly toward the cuttlefish.

Two experiments were carried out: 1) The memory was queried based on each object’s expected value (“What”). 2) The memory was queried based on each region’s expected value (“Where”). The parameters for this scenario are given in Table 2.

**Table 2.**
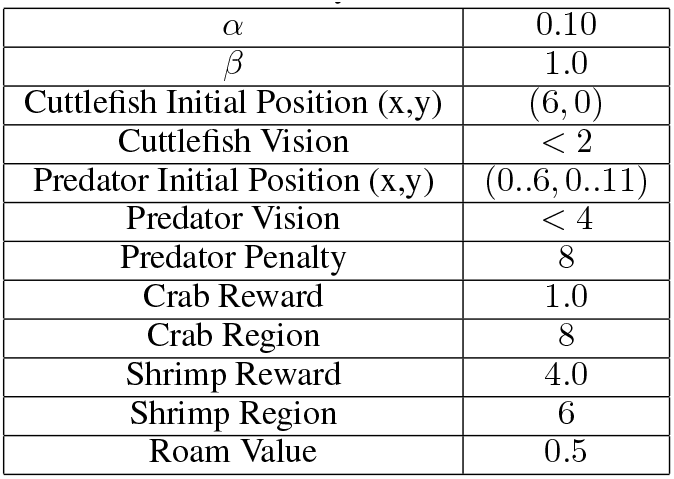
Predator-Prey Scenario Parameters.

|  |  |
| --- | --- |
| $\alpha$ | 0.10 |
| $\beta$ | 1.0 |
| Cuttlefish Initial Position (x,y) | (6, 0) |
| Cuttlefish Vision | < 2 |
| Predator Initial Position (x,y) | (0..6, 0..11) |
| Predator Vision | < 4 |
| Predator Penalty | 8 |
| Crab Reward | 1.0 |
| Crab Region | 8 |
| Shrimp Reward | 4.0 |
| Shrimp Region | 6 |
| Roam Value | 0.5 |

**What Query Experiment** At the start of every hour: 1) Values were acquired with a “what” query for each object (crab, shrimp) at the current time (see Equation 1). In this experiment, the total value summed over all locations for each object. 2) An action was chosen with the Softmax function (see Equation 4). 3) If the action was to *hunt*, an additional query then retrieves the region where the chosen object has its maximum value. The agent then proceeded to the center of the selected region. It is assumed that the cuttlefish agent knows how to reach this location. 4) If the action is to *roam*, then the cuttlefish agent moves a step in one of 8 cardinal directions. The direction is chosen randomly. 5) For all 3 actions, if a shrimp or crab are available at this time, and the cuttlefish agent is less than 2 steps from the object, the value is updated by Equation 2.

**Where Query Experiment** At the start of every hour: 1) Values were acquired with a “where” query for each region (see Equation 1). In this experiment, the total value was summed over all locations within each region. 2) An action for going to a region was chosen with the Softmax function (see Equation 4). 3) The agent heads from its initial position to the center of the chosen region. 4) If the action was to *roam*, then the cuttlefish agent moves randomly a step in one of 8 cardinal directions. The direction is chosen randomly. 4) For all actions except *hide*, if a shrimp or crab are available at this time, and the cuttlefish agent is less than 2 steps from the object, the value is updated by Equation 2.

## 3 Results

### 3.1 Episodic Like Memory Simulations

Episodic-like memory was shown in studies in which the cuttlefish had to remember where crab and shrimp were located, and when the preferred food (i.e., shrimp) was available [17, 31]. To show how the present episodic memory model could support such behavior, the basic idea of the study was replicated (see Figure 2). The cuttlefish agent roamed its environment until it either found a prey or 100 time steps, which corresponded to an hour, had passed. The simulation lasted 100 days and there were 3 hours in each day. Because the cuttlefish agents made random movements sometimes, the simulation was run 100 times, with each run lasting 100 days. In the first phase of the experiment, both crab and shrimp are available all day (3 hours). After 50 days, phase 2 begins where the crab is still available all 3 hours, but the shrimp is only available during the 3^rd^ hour.

**Figure 2.**
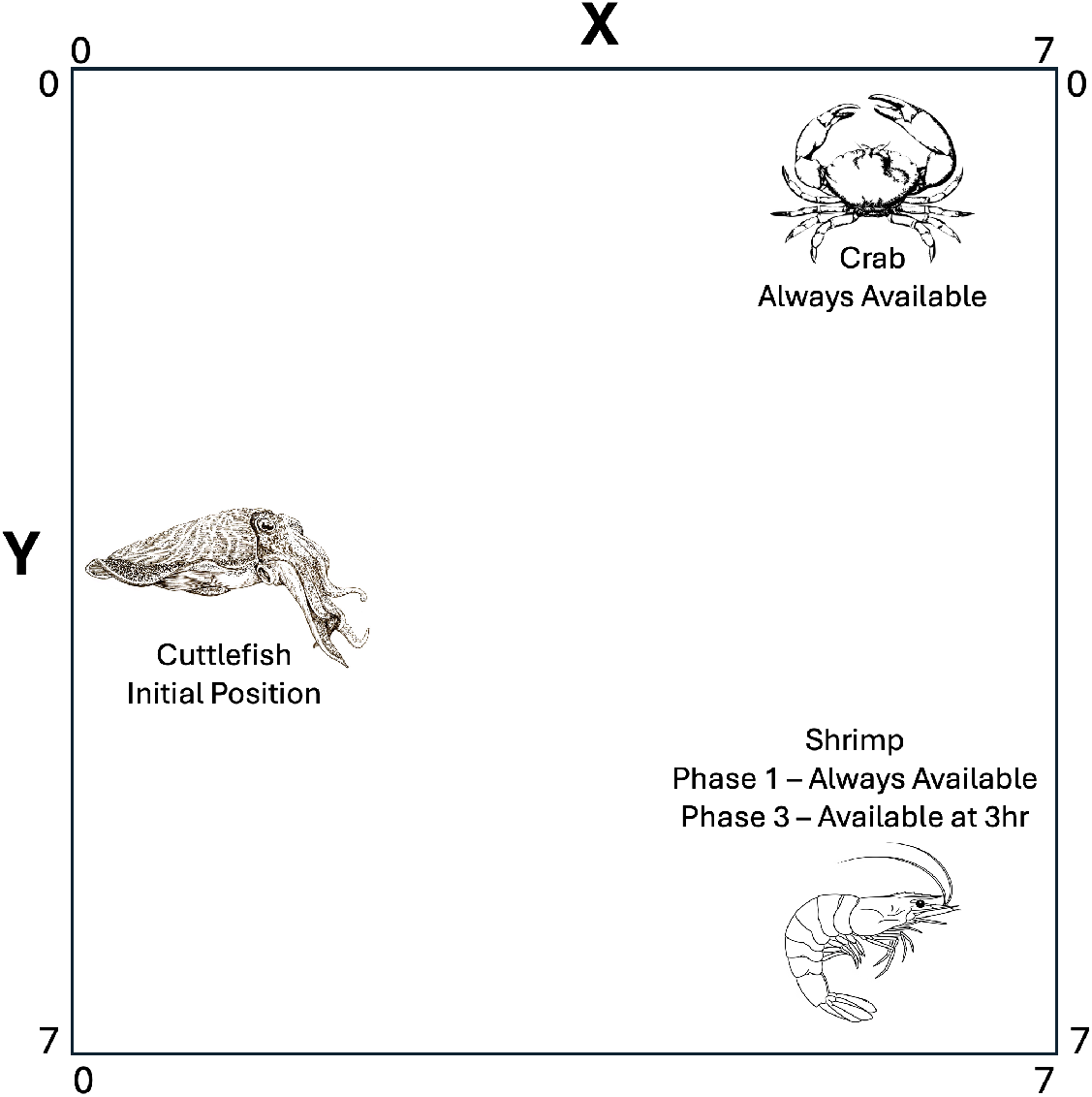
Episodic-Like Memory Environment. At the start of each hour, the cuttlefish is placed in the middle of the left side of the environment (x=0, y=4), and a crab is placed in the upper right region (x=7, y=0). The crab is available every hour. The shrimp’s location is in the bottom right region (x=7,y=7). During phase 1, the shrimp is available every hour. During phase 2, the shrimp is only available during hour 3. The cuttlefish can move freely within the environment, shrimp and crab remain stationary.

**Figure 3.**
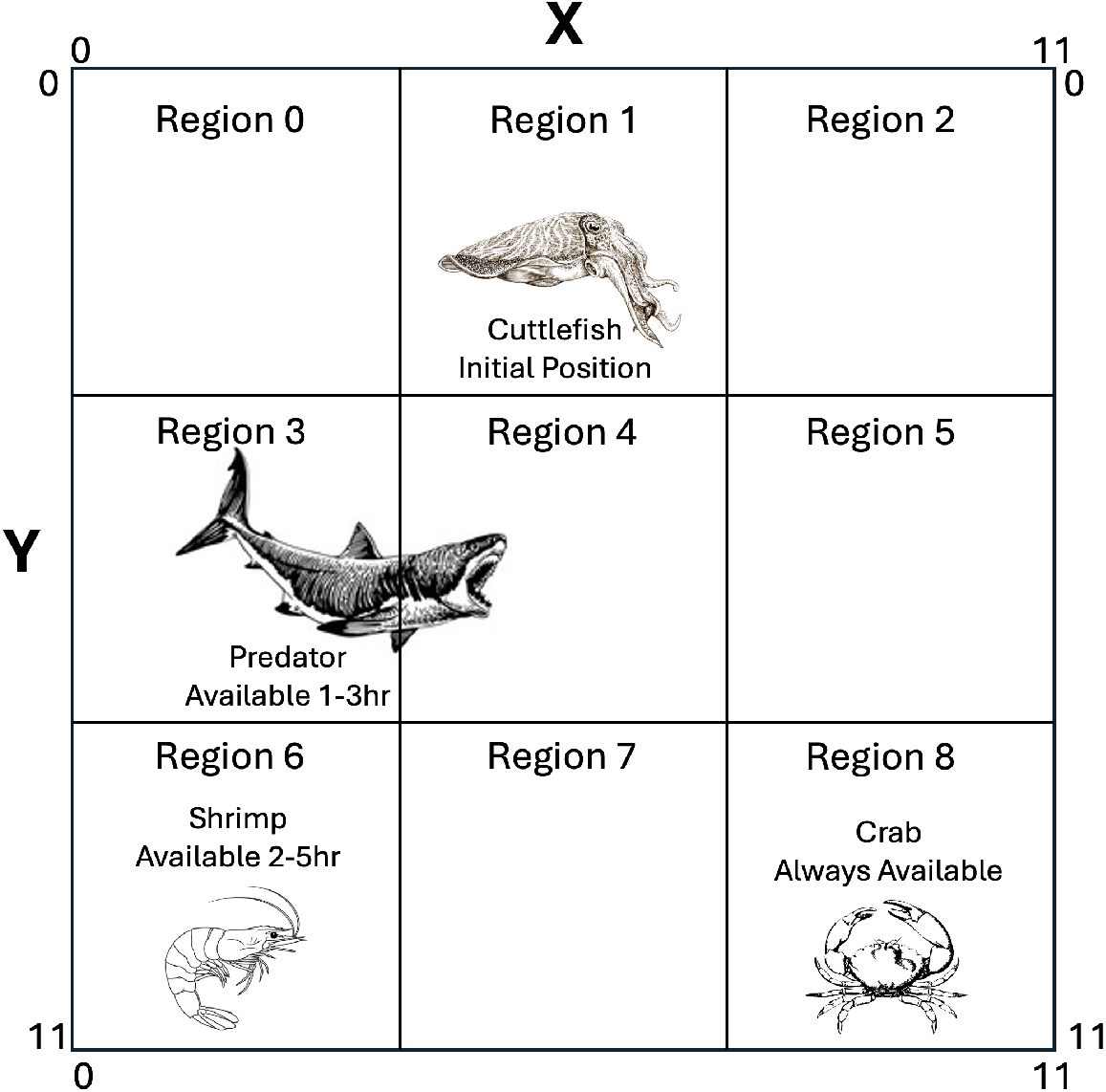
Predator-Prey Environment. The simulation environment is divided into 9 regions. Every hour, the cuttlefish is placed at the top (x=6, y=0) of region 1 and can move freely between regions. Shrimp are available from hour 2 to hour 5 in region 6. Crab are available all day (hour 0 to hour 5) in region 8. The predator is in the environment from hour 1 to hour 3. Every hour when available, the shrimp and crab are placed anywhere in their respective regions and stay stationary. Every hour when present, the predator is placed somewhere in the left side (X < 6) of the environment. The predator either moves randomly or approaches a cuttlefish if it is within the predator’s vision.

The cuttlefish agents choices in both phases were consistent with those reported in [17]. In phase 1, the cuttlefish agent showed a clear preference for shrimp (see Figure 4A.). In phase 2, the cuttlefish agent went to the crab location when there were 1 hour delays, and chose to forage in the shrimp location when there were 3 hour delays (see Figure 4B). These results show that the present memory model can support the acquisition and recall of “what”, “when”, and “where” information that is a feature of episodic-like memory.

**Figure 4.**
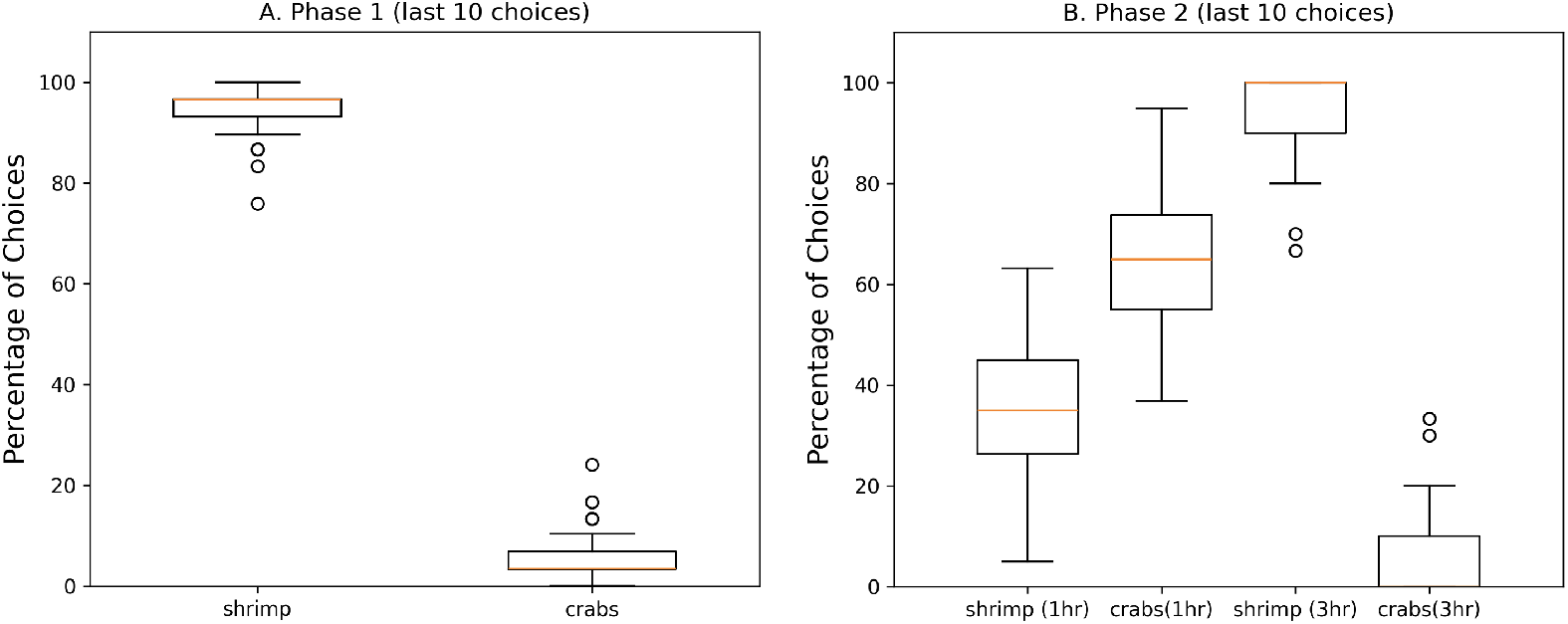
Episodic Like Memory. The percentage of choices after different delays are shown in the boxplots. The last 10 choices in each phase are shown for the 100 simulation runs. **A. Phase 1.** Crab and shrimp are available every hour. Phase 1 last the first 50 days. **B. Phase 2**. Crab are available after 1 hour delays. Shrimp are available 3 hour delays. Phase 2 lasts the second 50 days. The red line in the boxplot is the median of the 100 simulation runs, the box extends from the first quartile to the third quartile, the whiskers extend 1.5x beyond the quartiles. Circles denote outliers. Note that in phase 2, the median for crabs(3hr) is zero.

### 3.2 Predator-Prey Simulations

The Predator-Prey simulations further tested the episodic memory model by introducing a predator that appears a few hours each simulation day. The agent now must remember when and where prey types are available but also take into account when and where a predator might show up. As in the previous experiment, shrimp were preferred over crabs. Shrimp availability has more overlap with when and where the predator appears (see Figure 3). Note that the agent can only eat one prey per hour, or be eaten once per hour. After any of these events occur, the agent takes no further action until the next hour. Because the cuttlefish and predator agents have randomness to their movements, the simulations were run 100 times.

#### 3.2.1 What-When Query Experiments

In these experiments, the memory is first queried to get the expected values of the predator and each prey at the current hour. These values are turned into an action vector to *roam, hunt crab, hunt shrimp*, or *hide*. Figure 5 shows how the cuttlefish agent learned over time to avoid the predator, and eat the preferred prey. By the 100^th^ day, agent ate more shrimp than crabs and was rarely caught by the predator.

**Figure 5.**
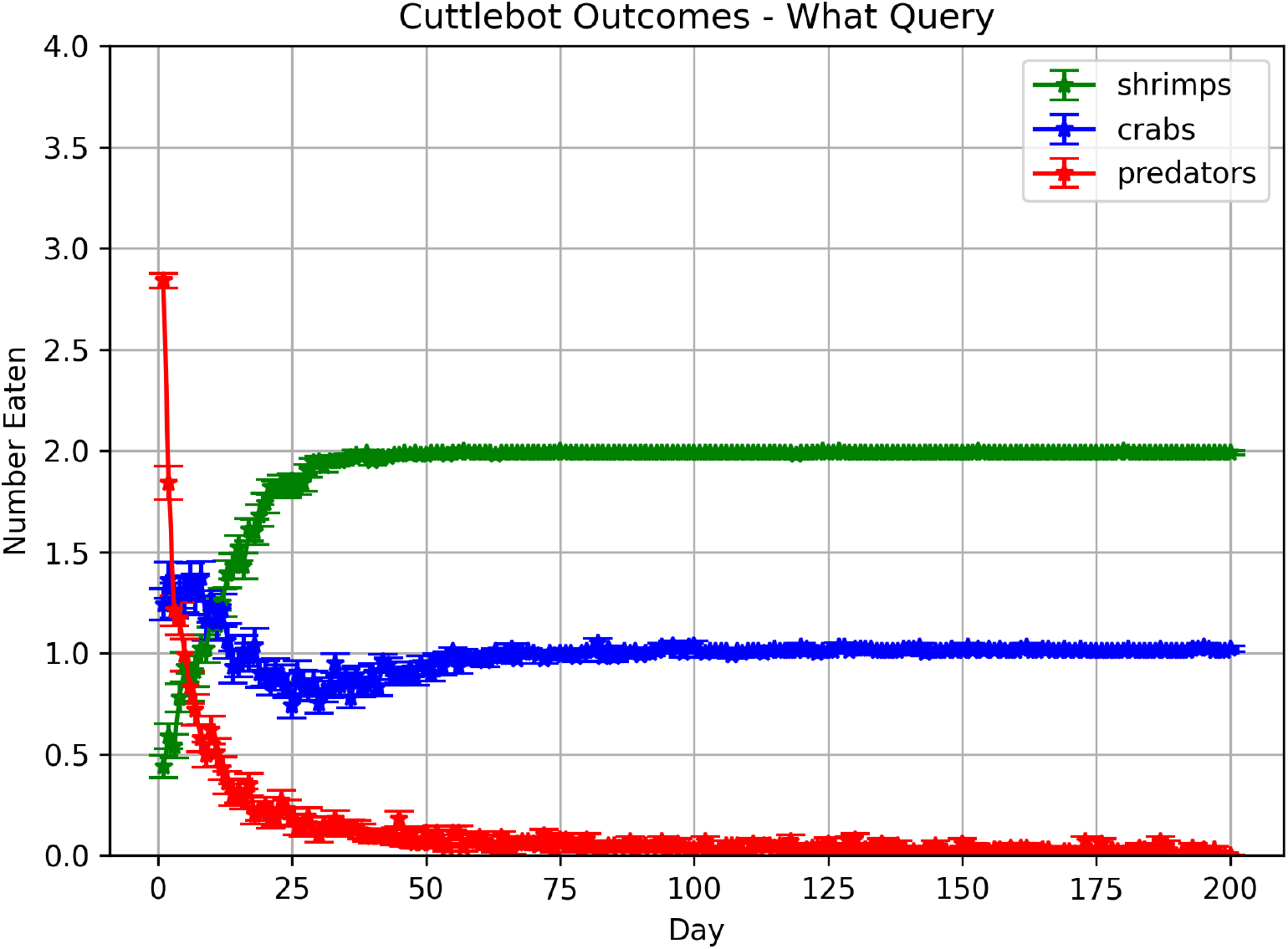
Outcomes in predatory-prey scenarios with “What-When” queries. Each point denotes the mean number of times the cuttlefish agent ate a specific prey or was eaten by the predator. The error bars denote the standard error.

An examination of the actions taken per hour revealed that the cuttlefish agent learned to opportunistically hunt crab and shrimp during the hours that the environment was predator free. In the first 20 days, the cuttlefish agent *roamed* often, which increased predation risk but also facilitated exploration of the environment (see Figure 6 A). By the last 20 days, the cuttlefish agent learned what objects to seek, when to seek them, and where to find them (see Figure 6 B).

**Figure 6.**
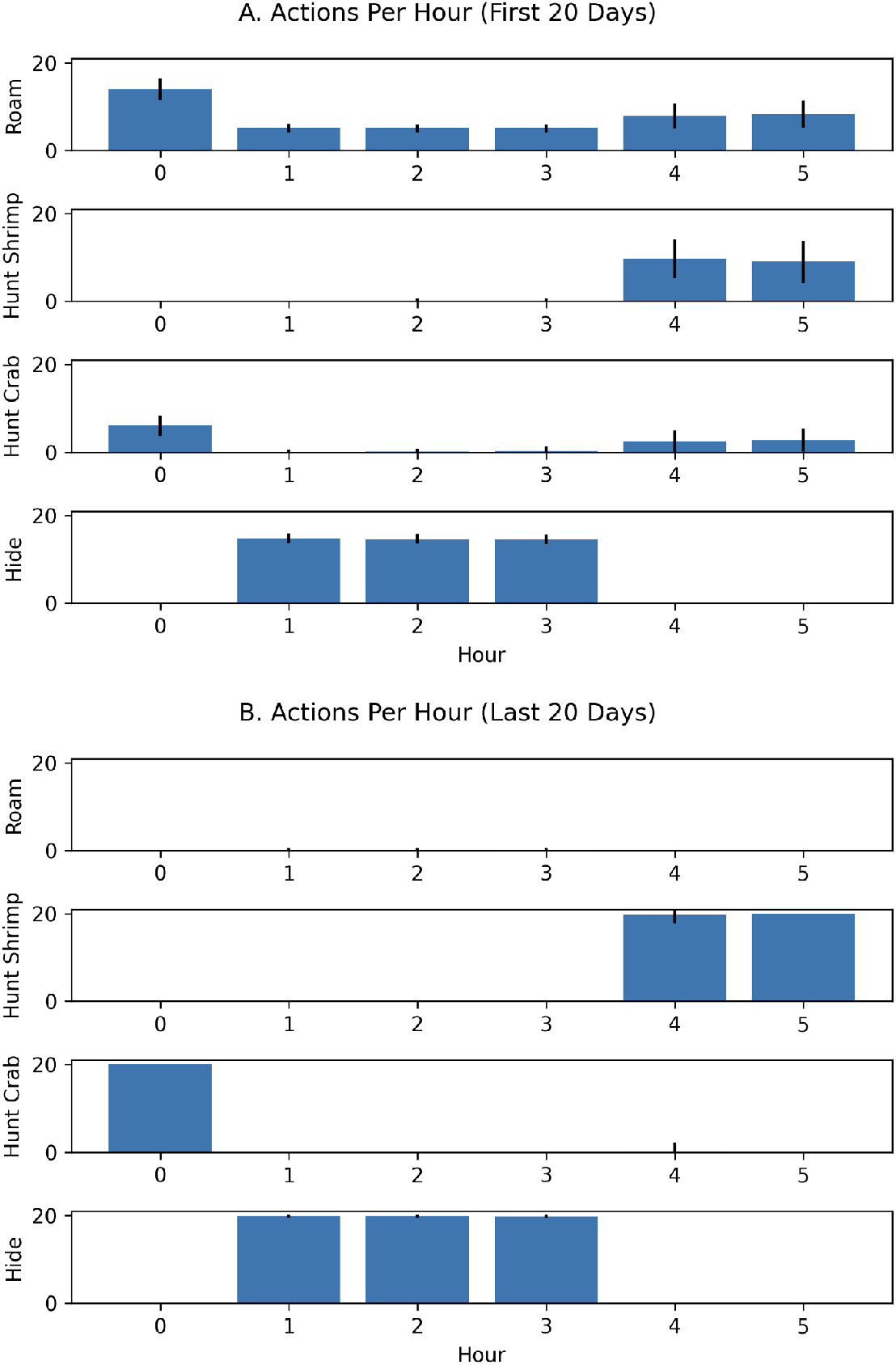
Actions per Hour with “What-When” queries. The roam, hunt or hide that the agent chose per hour is shown for early and late trials. **A.** Actions chosen on the first 20 days. **B**. Actions chosen on the last 20 days. Bars denote the mean actions taken per 100 runs and the error bars denote the standard deviation.

#### 3.2.2 When-Where Query Experiments

In these experiments, the memory was queried to get the expected values of each region at the current hour. These values were turned into an action vector to move towards a region in the environment. Figure 7 shows how the cuttlefish agent learned over time. In contrast to the “What-When” query experiments, more crabs are eaten than shrimp. Although the cuttlefish agent learned to somewhat avoid the predator, it was still occasionally eaten. The reason for this is that the path taken to the crab was, for the most part, predator-free. It appears that the agent opportunistically hunted crab which was a safer choice.

**Figure 7.**
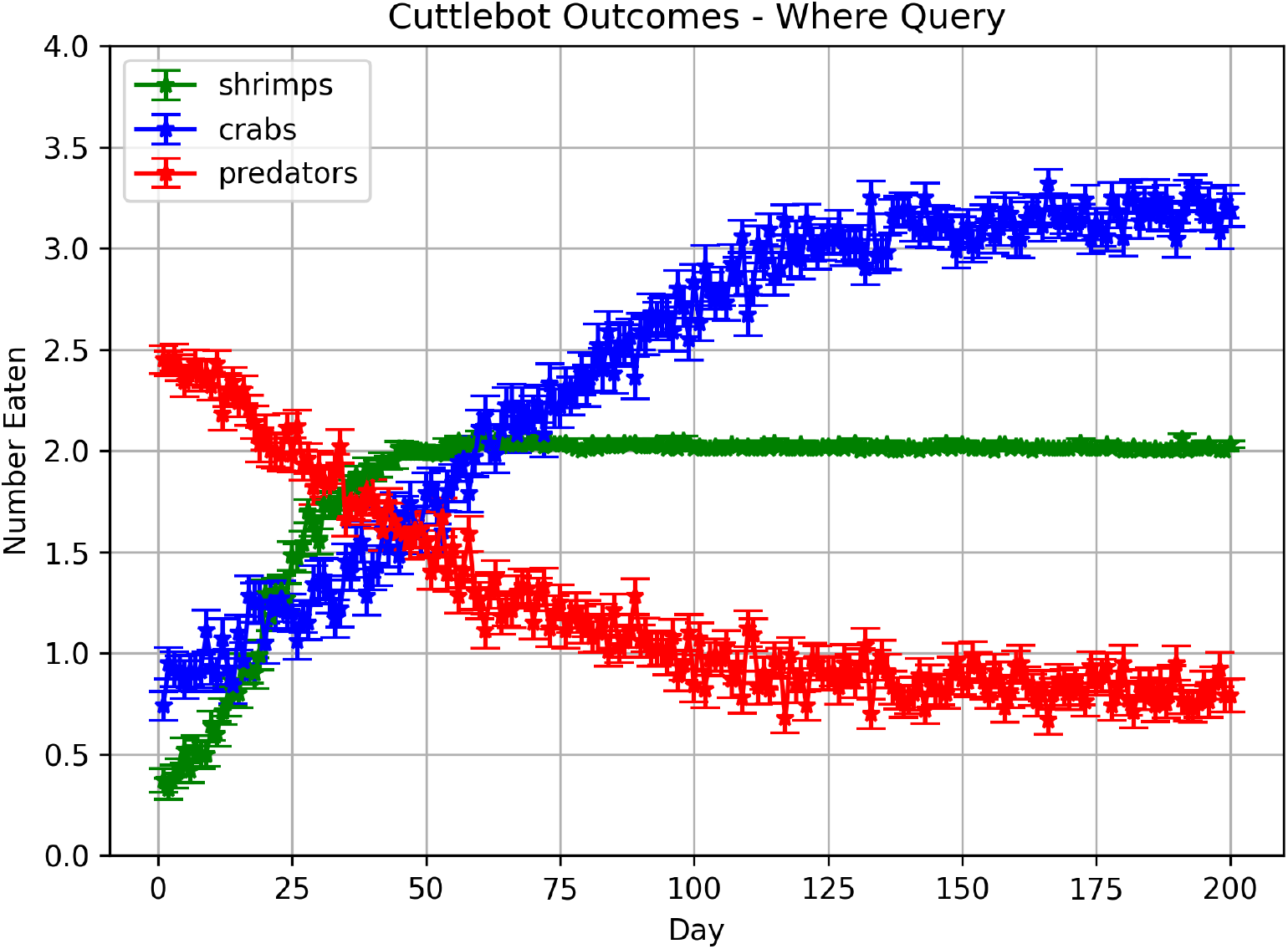
Outcomes in predatory-prey scenarios with “When-Where” queries. Each point denotes the mean number of times the cuttlefish ate a specific prey or was eaten by the predator. The error bars denote the standard error.

An examination of the actions taken per hour in these experiments revealed that the cuttlefish agent learned the environment in the first 20 days by sampling all regions (see Figure 8 A). By the last 20 days, the cuttlefish only explored the two regions where prey are found (see Regions 6 and 8 in Figure 8 B). In region 6, where shrimp are found, the cuttlefish agent primarily went there during hours 5 and 6 when it was predator-free. Yet it still occasionally looked for shrimp in hours 2 and 3 when shrimp were available but the predator was present. During hours 0 through 4, the cuttlefish agent opportunistically sought crab in region 8, even those times when the predator was present. However, when shrimp were available risk-free (hours 4 and 5), the cuttlefish agent only explored region 6.

**Figure 8.**
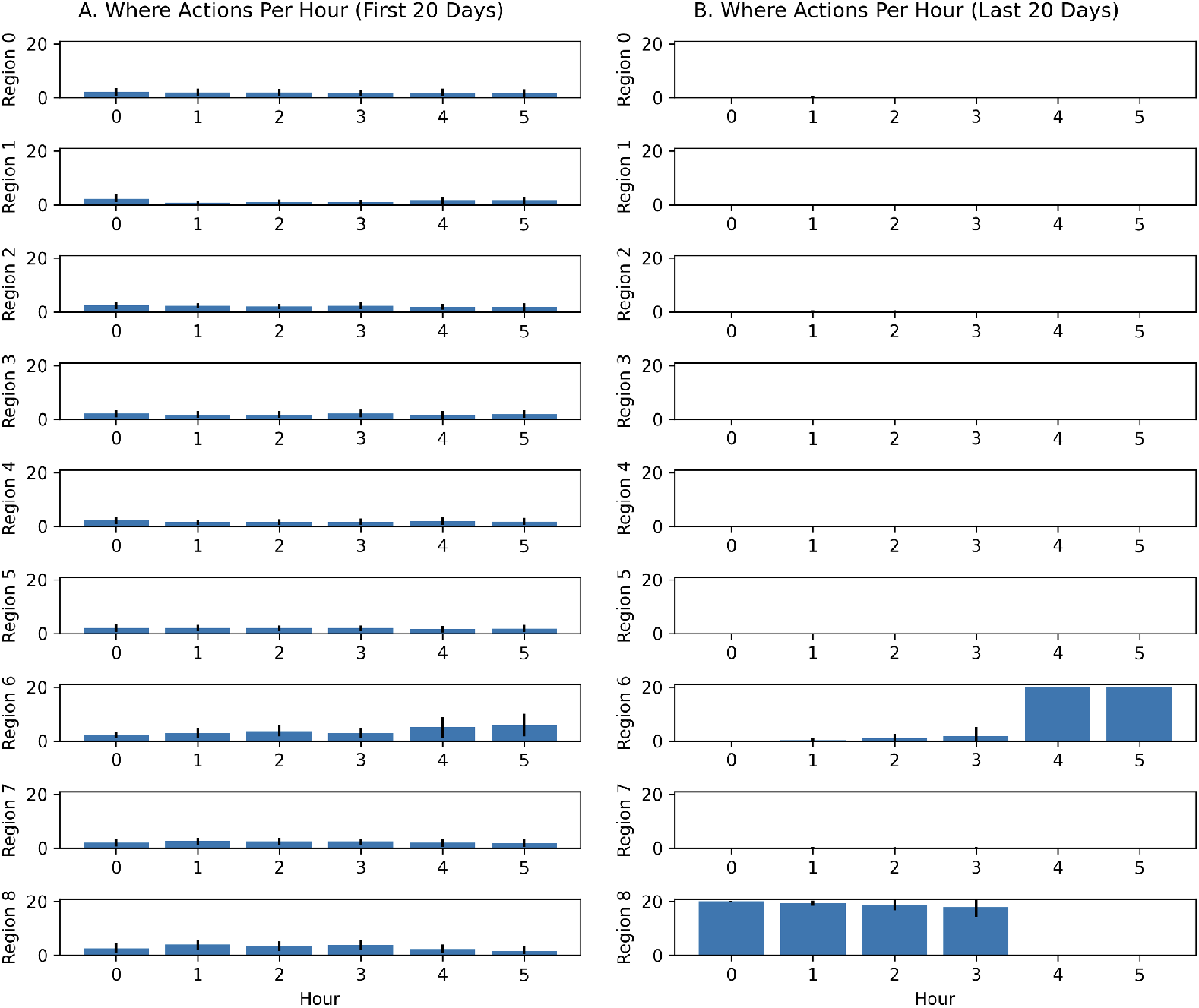
Actions per Hour with “When-Where” queries. The region that the agent chose per hour is shown for early and late trials. **A.** Actions chosen on the first 20 days. **B**. Actions chosen on the last 20 days. Bars denote the mean actions taken per 100 runs and the error bars denote the standard deviation.

### 3.3 Mental Time Travel Through Memory Queries

Mental time travel is the ability to reason through time, and it is a hallmark of episodic memory. The episodic memory model has sufficient information to show this capability. Because queries can reconstruct past or anticipate future states, the model supports a rudimentary form of mental time travel. For instance, similar to animal experiments of episodic-like memory, the behavior shown in Section 3.2 hints at mental time travel being used, but lacks the declarative report.

Since we have access to the cuttlefish agent’s memory matrix, we can query the memory after its experience in the environment. Figure 9 shows the results of such queries at different times in the experiment. To create these charts, we made “what”, “when”, “where” queries to the memory after a typical simulation run. If it the time was hour 1, one could imagine traveling forward in memory time until hours 4 and 5 where the shrimp could be eaten without the risk of encountering a predator. Or traveling back in time to hour 0 for a risk-free crab breakfast.

**Figure 9.**
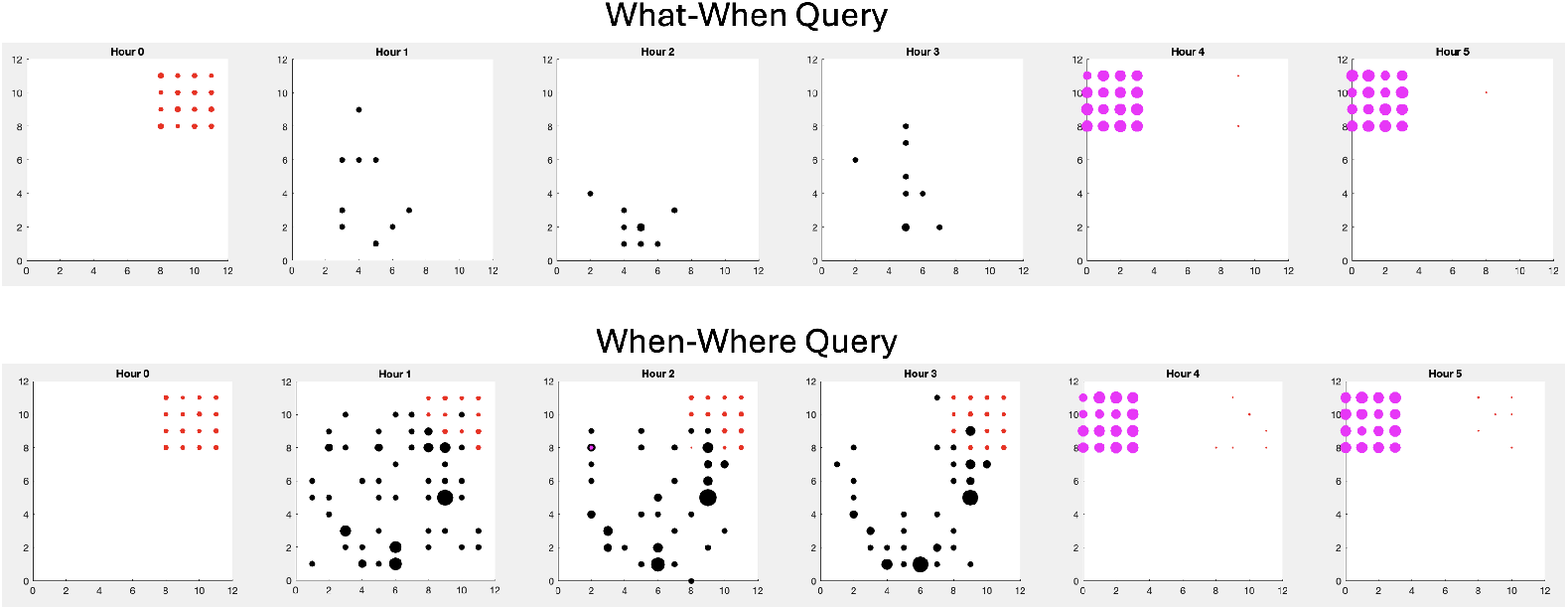
What, When, Where queries to memory. The scatter plots show the expected values of each object, at each location in the environment. The panels from left to right show the result at each hour of the simulation day. The magenta markers denote shrimp encounters, the red markers denote crab encounters, and the black markers denote predator encounters. The size of the marker is proportional to the expected value. The charts show the result after a typical simulation run. **Top.** Result of the “What-When” query experiments. **Bottom**. Result of the “When-Where” query experiments.

Figure 9 also illustrates how the type of query can shape the memory. The top chart in Figure 9 shows the result of the “What-When” query experiment. In this case, the agent took less risks by hunting prey when the predator was not present. The bottom chart of Figure 9 shows that “When-Where” queries led to opportunistic but risky crab hunting in hours 1 through 3. Note the larger expected value of the predator and the locations of the predator cover more environment. This is because the cuttlefish agent was not hidden (i.e., camouflaged) when hunting crab and may have encountered a predator.

## 4 Discussion

Episodic-like memory allows animals to contextualize past events in order to guide future actions. An episodic memory contains what happened, when it happened, and where it happened. This conjunction of what, when, and where has now been observed in a wide range of organisms including primates, birds, and cephalopods [8]. It has been argued that only humans have true episodic memory, where they can carry out mental time travel and reflect on their past. However, it is an open question whether this is because we share a common language with humans that allows us to report and probe memories in ways that we cannot with other animals. There is growing consensus that many animals have “dimensions of consciousness”, which not only include such self-reflection, but also other cognitive attributes [5].

In the present work, we introduced a parsimonious memory model that can replicate the episodic-like memory shown in cephalopod behavioral experiments [17, 31], as well as other scenarios requiring “what”, “when”, and “where” conjunctions to recall episodic events. The model takes inspiration from the hippocampal indexing theory [35, 36] in that combinations of “what”, “when”, and “where” indices can be used to store and recall memories. The cephalopod’s vertical lobe, which is important for learning and memory, has anatomical similarities to the hippocampus [32, 33]. Therefore, the model presented here may be applicable to some birds and mammals.

In simulations, our episodic memory model: 1) Replicated episodic-like memory experiments in cuttlefish [17]. The simulated cuttlefish agent remembered that preferred food could be found at a certain location after longer time delays than a non-preferred food. 2) Demonstrated episodic-like memory in more complex simulations that required flexible behavior. A predator-prey scenario required the cuttlefish agent to remember not only when and where preferred and non-preferred food could be found, but also had to remember when and where a predator might be encountered. The agent successfully caught preferred food when the predator was not present, and opportunistically caught non-preferred food by avoiding the predator’s typical locations. 3) Showed that how the memory was queried could affect the agent’s behavior. If the query was for what objects could be found when, the agent demonstrated risk-averse hunting. If the query was for when and where objects could be found, the agent opportunistically hunted prey while risking encounters with the predator. In both cases, the cuttlefish agent successfully learned to avoid predators and hunt prey. 4) Demonstrated the ability to travel forward and backward through time. Queries of the memory showed the expected value landscape over time. Although not shown here, a reasoning algorithm could autonomously play out imagined scenarios to choose the appropriate time and place to act.

### 4.1 Model Assumptions

The memory model presented here was purposefully made simple to show how such a memory structure could support episodic-like memory. Therefore, we assumed the agent had *a priori* abilities that were not necessarily part of the episodic memory, but required for the agent to carry out its behavior.

We assumed that once the agent remembered an object’s location, it knew how to get near there. We loosened this assumption by dividing the environment into regions and once in a region the agent had to search for its prey. If path planning is an important feature of future versions, then the present model could readily incorporate other sequence learning algorithms to plan paths to locations, such as reinforcement learning, successor representations or recurrent neural networks [4, 10, 12, 34].

Another assumption was that the agent had a sense of time passage. Although it is clear in humans and other organisms that they have a sense of time at multiple timescales, it is not agreed upon how the brain implements an internal clock. Time cells have been reported in the rodent and human hippocampus [9, 40]. Others have suggested that the basal ganglia and its interactions with the cortex can keep track of time [22]. For now, we assume that there is a signal that denotes the passage of time and how that timekeeper is implemented is an open issue.

It has been proposed that neurogenesis in the dentate gyrus, which is a hippocampal subfield, could register memories in time [1, 2]. The “when” component of an episodic memory could be encoded by a neuron’s birth date. Interestingly, the octopus nervous system grows and shrinks dramatically during its lifetime (W-S. Chung and F. Cortesi, personal communication). This neurogenesis, as the cephalopod’s brain grows with experience, fits with the dentate gyrus time encoding idea.

### 4.2 Mental Time Travel

The simulations in Section 3.3 showed that the memory model could be queried forward or backward in time. Using this information would allow the agent to observe the memory landscape and decide to act now or delay action. Mental time travel, which involves imagining past memories and looking to the future, is a hallmark of episodic memory. While it cannot be said that the present model has this capability, it could be expanded to support this idea of mental time travel. It would require the addition of reflection and reasoning over the stored memories. In [26], this was achieved by implementing a large language model with a memory store and a reflection module. Such additional models could be integrated with the present episodic memory structure.

### 4.3 Comparison to Other Models

The hippocampus and the surrounding medial temporal lobe have inspired numerous memory models. These models typically focus on the spatial navigation that is a major cornerstone of hippocampal research [4, 12, 13]. Hippocampus inspired models such as the Byrne, Becker, Burgess (BBB) [6], clone-structured cognitive graph (CSCG) [13] and the Tolman-Eichenbaum Machine (TEM) [41] support memory with different relational structures. BBB suggests that the parietal and retrosplenial cortices support transformations between egocentric and allocentric representations. It uses the head direction system to rotate these representations and the hippocampus to access memories [6]. CSCG uses graphical models to represent memory by creating different clones of observations for different contexts to resolve spatial ambiguity [13]. CSCG focuses on sequential learning with a graphical model that ties together observations with hidden states. When a new memory is acquired or an established memory deviates, a clone of the structure is created. These may be linked together through the graph structure. TEM proposes that entorhinal cortex cells form a structural knowledge basis while hippocampal cells link this basis with sensory representations, allowing generalization across environments with similar structures. The TEM focuses on relationships that form memories. In their work, they use spatial navigation, family relations, and semantic relations as examples. These models are specific to the mammalian hippocampus and may not be applicable to other biological memory structures like the cephalopod vertical lobe. Moreover, these models require complex computations and data structures to perform memory functions. They also don’t address the conjunctions, especially when events occurred, necessary for episodic memory.

A major goal of the present work was to show episodic-like memory in a simple, yet biologically plausible data structure. The mammalian hippocampus, which appears to be necessary for episodic memory, receives processed sensory input from numerous brain regions [29, 30, 37]. Evidence suggests that the hippocampus accesses memories with conjunctions of these inputs [19, 18]. Thus, the choice of a three-dimensional matrix with dimensions denoting the what, when and where aspects of a memory, although overly simple, is justified. The anatomy of the cephalopod vertical lobe, which like the hippocampus receives multiple sensory input streams [32, 33], further justifies this memory model.

The present memory model has similarities to content addressable memories like Hopfield networks [15]. In a Hopfield net a complete memory, which is stored in a matrix-like structure, can be retrieved with partial cues. Modern versions of Hopfield networks can create and access relational memories with queries [3, 21]. These network models are loosely based on biological memory structures. Like other models discussed, they are not focusing on conjunctions of what, when, and where.

### 4.4 Model Extensions

A shortcoming of the present model is that it doesn’t scale and it is only accessed across the three dimensions. The scaling issue could be addressed with sparse coding and quantized query keys as in [3]. The original hippocampal indexing theory suggested that a partial index into the hippocampus could yield index pointers to multiple cortical areas to bring forth a complete memory [35]. In computer memory circuitry, content addressable storage systems use a hash code and other methods to scale up these memory systems [23]. Such structures could readily be integrated into the present episodic memory model.

The behavior and environment in the present simulations are highly constrained. This was purposely done to replicate controlled experiments and to show how the present memory structure could support behavior in a more challenging scenario. As discussed, the present model could be extended to support path planning and a more plausible representation of time, which would make it’s behavior more complete.

The episodic memory model introduced here is a part of an ongoing project called CuttleBot. One of the first outcomes of this project was a biomemetic robot [28]. The highly constrained environment issue might be addressed if this memory model was embodied in an autonomous robot with active sensing. The field of neurorobotics connects the brain, body and environment [16]. The three main design principles for neurorobots are [20]: 1) they must react quickly and appropriately to events, 2) they must have the ability to learn and remember over their lifetimes, 3) they must weigh options that are crucial for survival. It is believed that following these design principles makes the robot’s behavior more realistic and successful. The latter two principles are addressed in this paper. The present model supports learning and memory, and the agent must weigh the tradeoff between foraging for food and becoming food itself. However, the first principle is grounded in the idea of embodied cognition, that is, body morphology and interaction with the real-world shapes behavior that responds quickly to events. By incorporating the present work into a robot, all three principles could be addressed.

### 4.5 Conclusion

In summary, we introduce a memory model that supports conjunctions of “what”, “when”, and “where” that are necessary for episodic-like memory. The data structure can be queried in any combination of these three dimensions, enabling recall of events in ways that parallel behavioral findings in cuttlefish. This suggests that such a structure could explain the episodic-like behavior observed across species. With some extensions it may also support mental time travel, which is thought to be a requirement for episodic memory. These findings highlight how foundational, biologically inspired memory structures can advance our understanding of cognition across species.

## Acknowledgments

The CuttleBot team was supported by the UC Irvine California Institute for Telecommunications and Information Technology (CALIT2) in collaboration with the UC Irvine Undergraduate Research Opportunities Program (UROP). JLK was supported in part by NIH award R01 NS135850-02. The authors would like to thank members of the CuttleBot team for many valuable discussions.

## Notes

https://github.com/jkrichma/EpisodicLikeMemoryModel.git

### Competing Interest Statement

The authors have declared no competing interest.

